## Supplemental Materials for "Prominin 1 and Tweety Homology 1 both induce extracellular vesicle formation"

***Supporting Information***

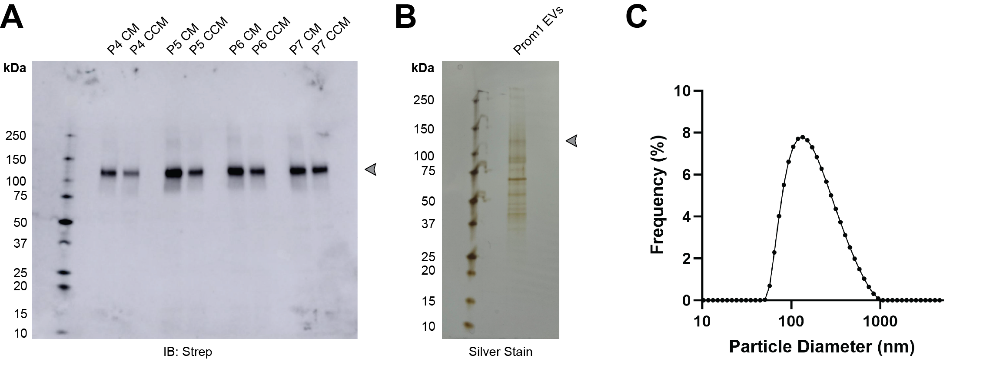

**Figure S1.** **(A)** Anti-Strep immunoblot of conditioned media (CM) or clarified conditioned media (CCM) from a stable polyclonal Expi293 cell line expressing lentiviral-transduced Prom1-Strep. Arrowhead indicates the expected position of glycosylated Prom1-Strep. P4, P5, P6, and P7 indicate the passage number of the suspension cell culture. **(B)** Total protein stain of purified Prom1-Strep EVs. Arrowhead indicates the expected position of glycosylated Prom1-Strep. **(C)** Dynamic light scattering (solution size) measurement of purified Prom1 EVs.

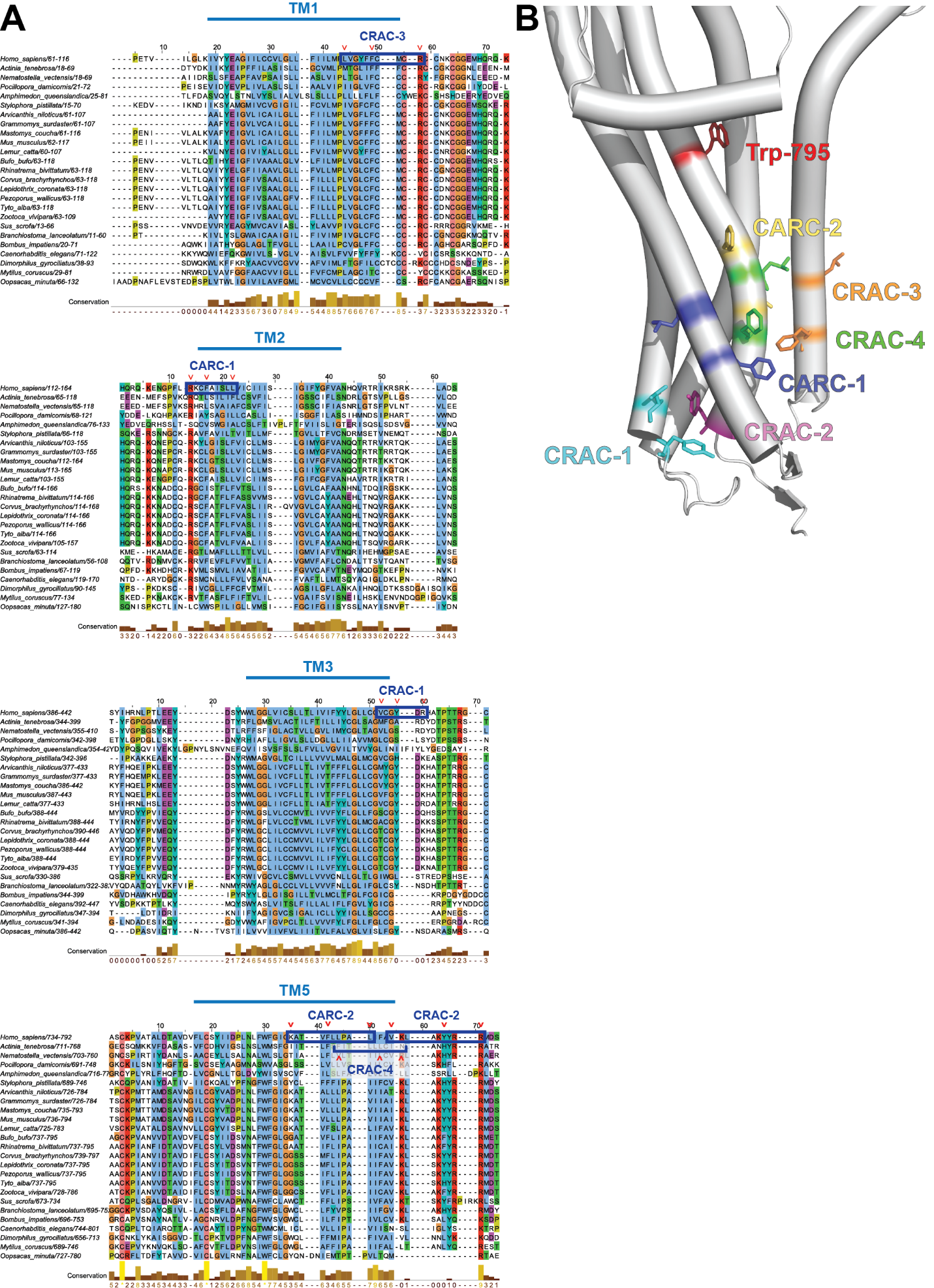

**Figure S2. (A)** Multiple sequence alignment of metazoan Prom1 focused on transmembrane domains 1, 2, 3, and 5, with human CRAC and CARC domains highlighted. Red carats indicate key charged, aromatic, or hydrophobic residues that define the CRAC and CARC domains. Alignment visualized using Jalview^1^. **(B)** Trp-795, CRAC-1, CRAC-2, CRAC-3, CRAC-4, CARC-1, and CARC-2 mutation sites superposed onto an AlphaFold2 model^2^ of the transmembrane domain of Prom1.

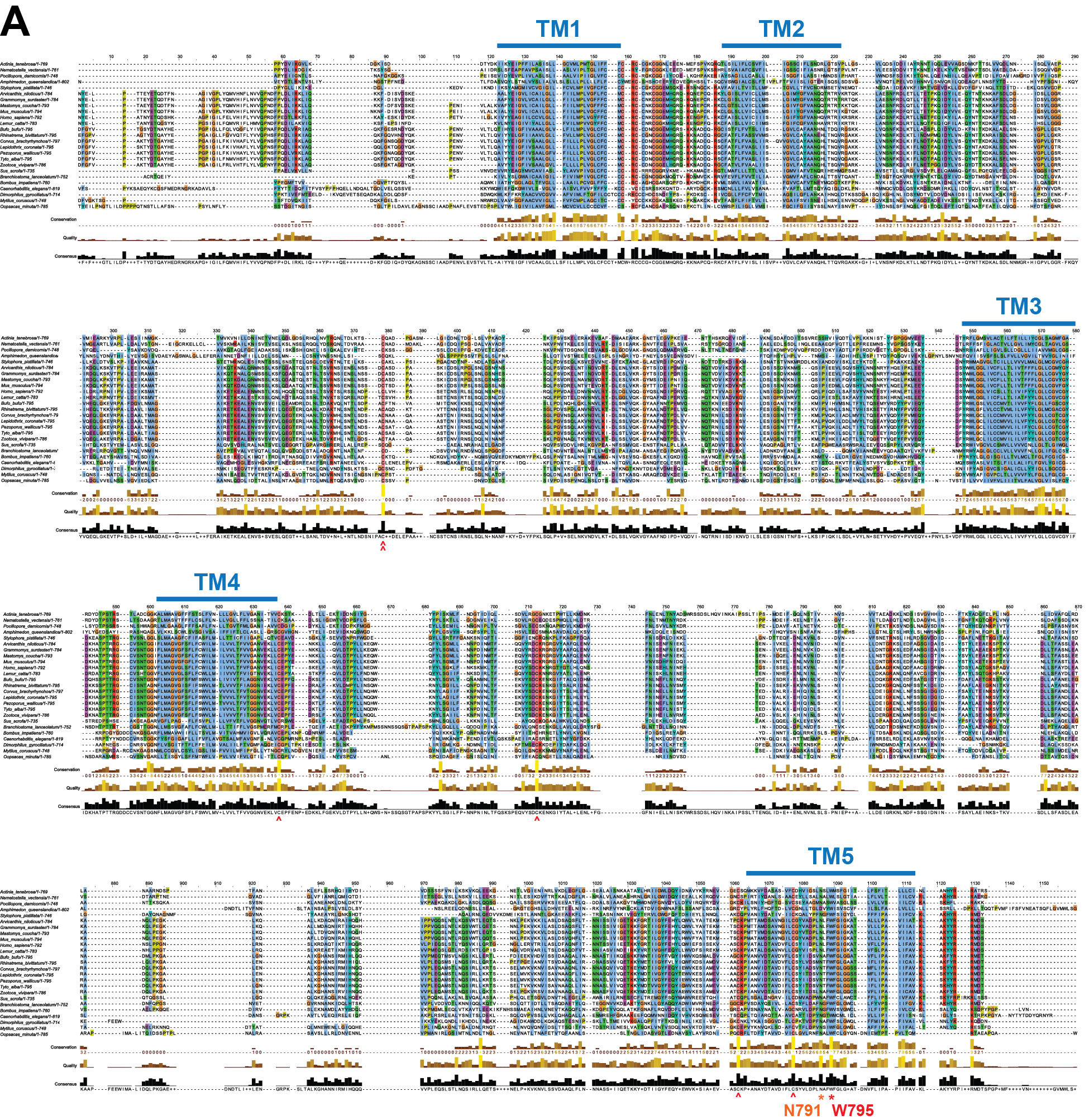

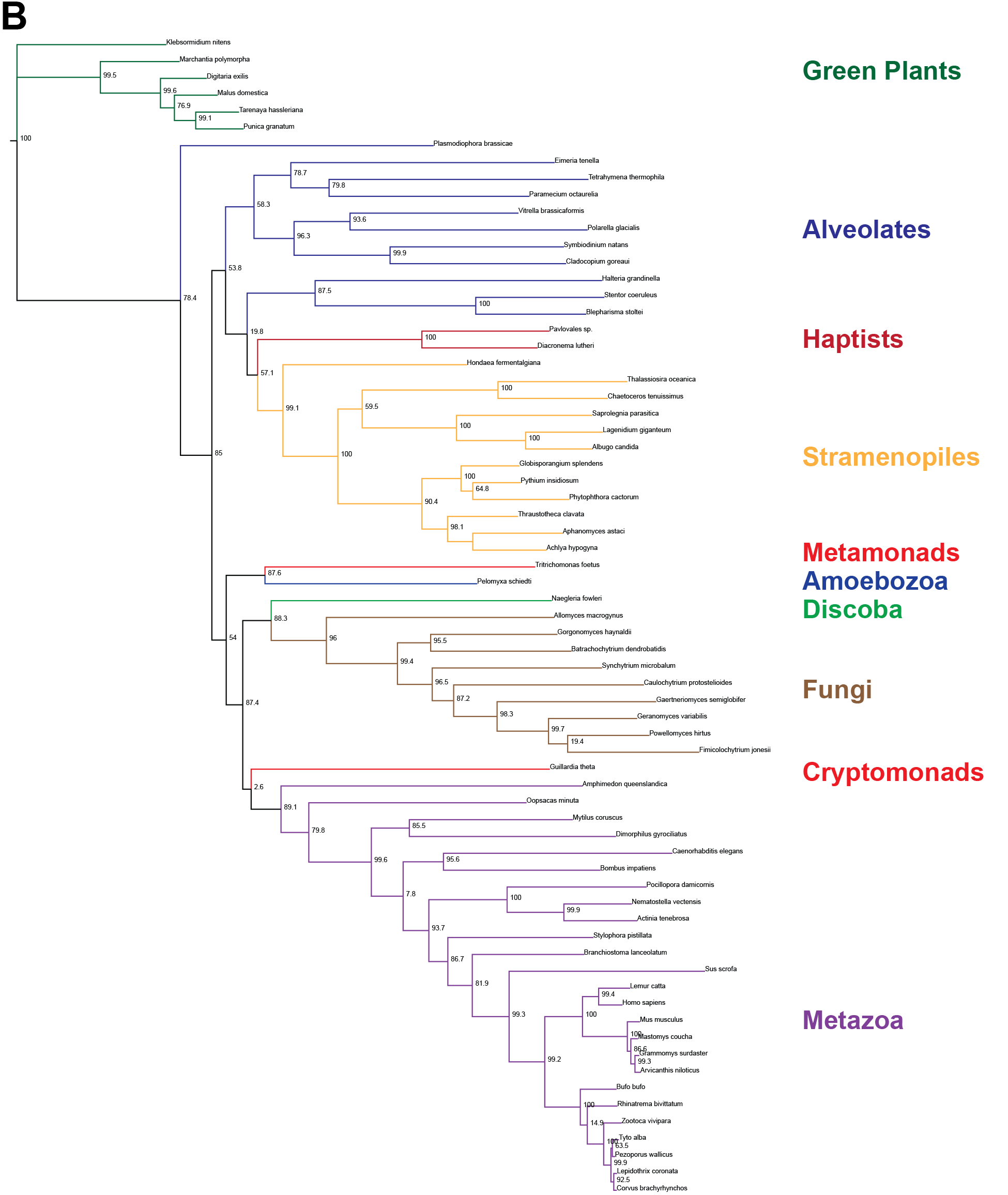

**Figure S3.** **(A)** Multiple-sequence alignment of prominin sequences from metazoa with the five transmembrane segments indicated. Red asterisk indicates perfectly conserved non-cysteine residues. Orange asterisk indicates less-than-perfectly conserved non-cysteine residues of interest. Red carat indicates cysteines predicted to form internal disulfides. Double red carat indicates cysteines that are not predicted to form internal disulfides. Alignment visualized using Jalview^1^. **(B)** Inferred phylogenetic relationships between putative prominin homologs identified across eukaryotes. Node labels indicate aLRT branch supports^3^. Tree visualized using IcyTree^4^.

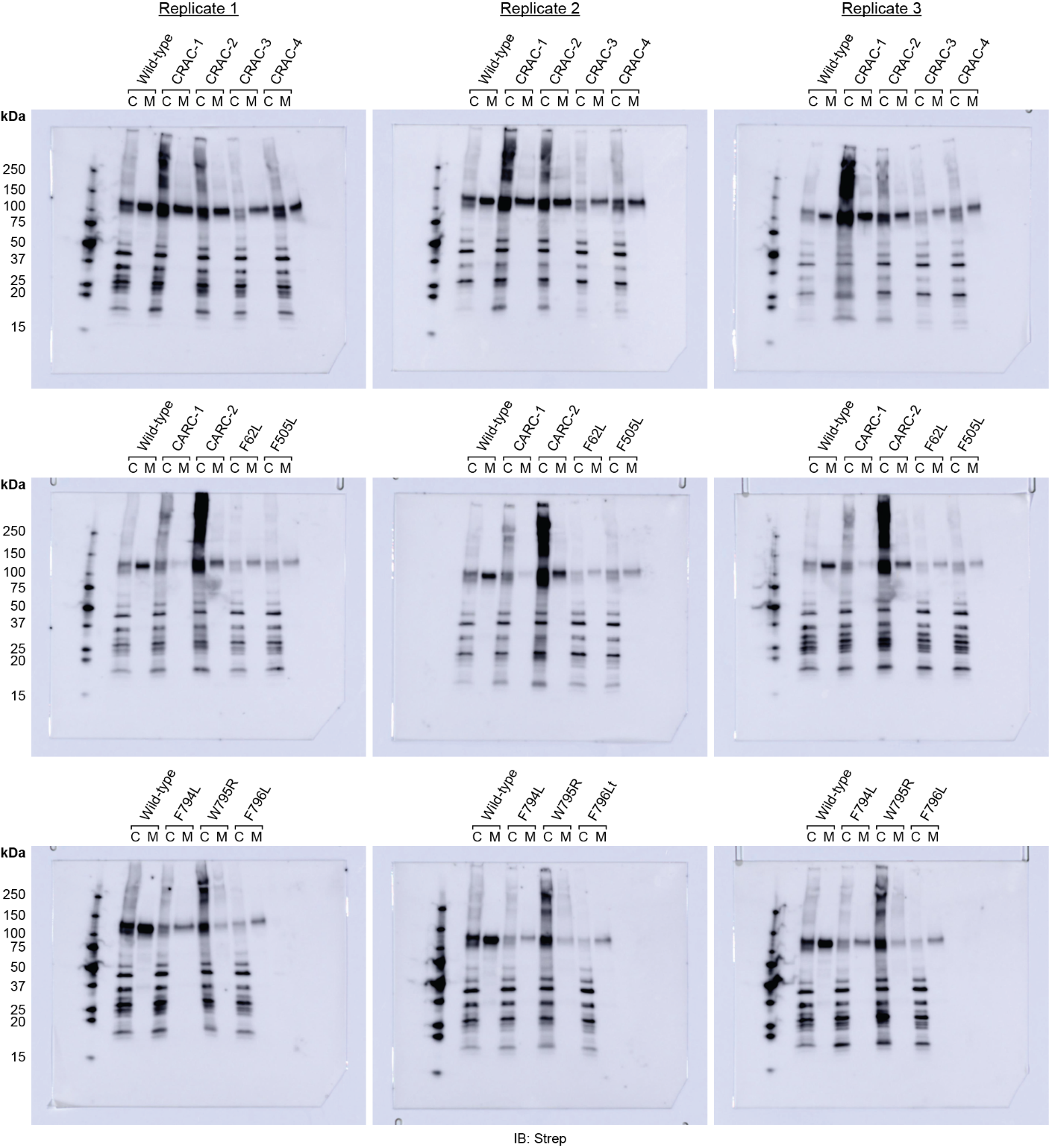

**Figure S4.** Anti-Strep immunoblots used for quantification of EV production by Prom1 mutants in Figure 2C. Lanes are labeled “C” for cells and “M” for clarified conditioned media.

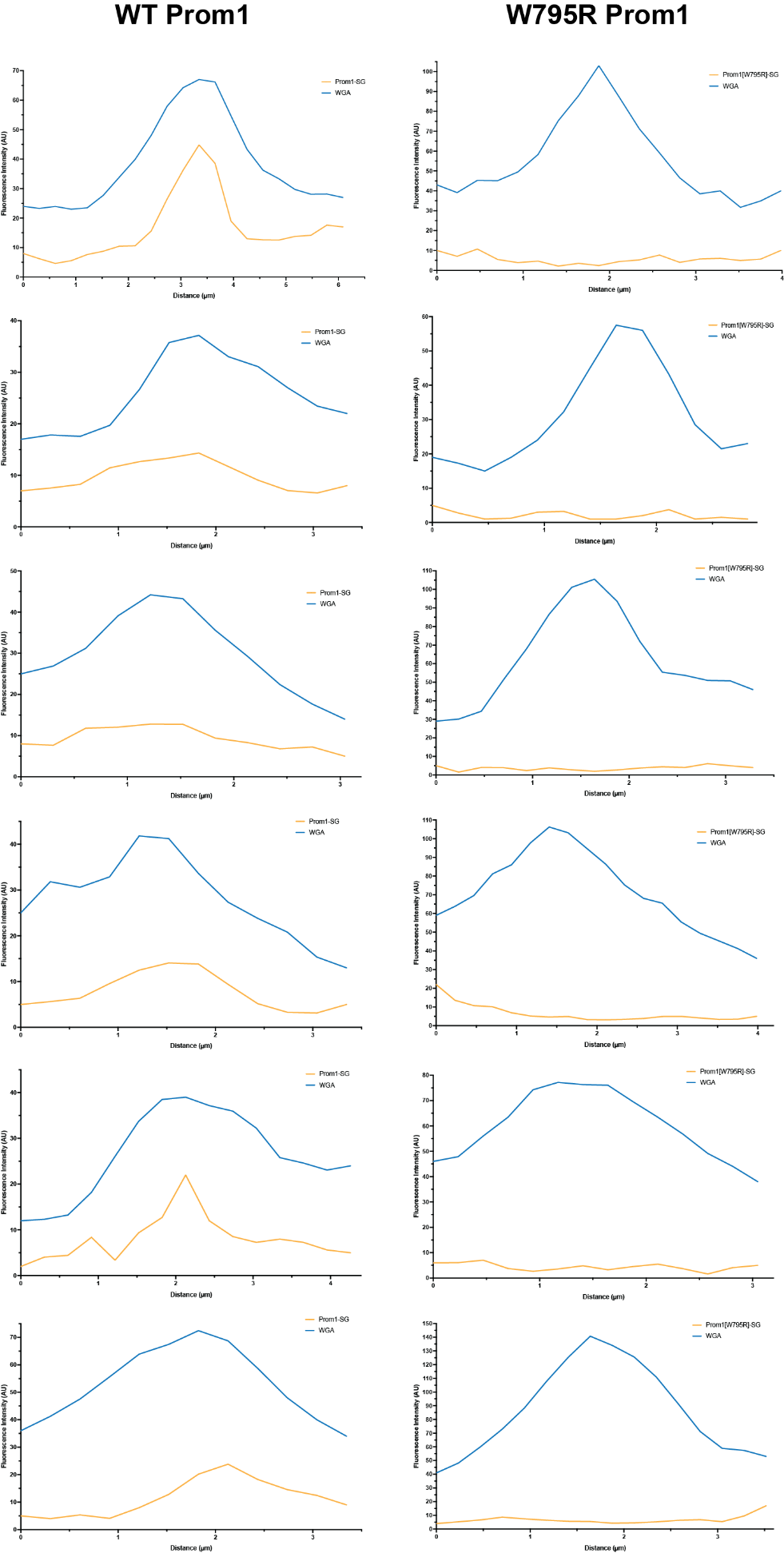

**Figure S5.** Line scan traces across cell junctions (n = 6) for plasma membrane (WGA) (blue) or Prom1-mStayGold fluorescence signal for WT (*left panels*) or W795R (*right panels*) Prom1 (yellow).

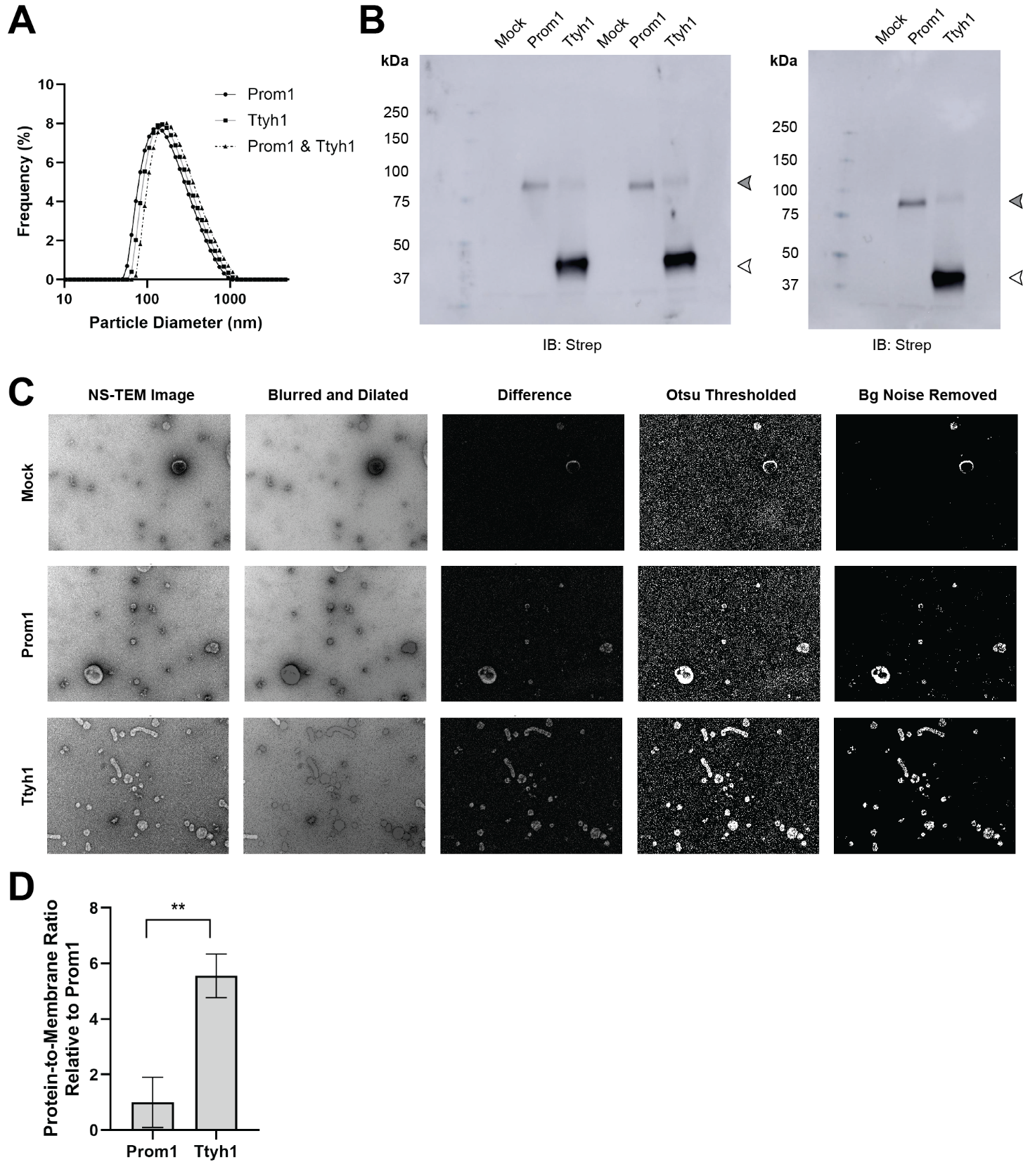

**Figure S6. (A)** Dynamic light scattering measurement of EV diameter for purified Prom1-Strep, Ttyh1-Strep, or Prom1-Strep + Ttyh1-Strep co-expression EVs. **(B)** Anti-Strep immunoblots used for relative quantification of Prom1 and Ttyh1 levels in purified EVs. Filled and empty arrowheads indicate the expected positions of Prom1 and Ttyh1, respectively. **(C)** Representative NS-TEM micrographs and image processing intermediates used for membrane area quantification. **(D)** Quantification of relative protein-to-membrane ratio for Prom1 and Ttyh1 EVs using data from Figures 3I and 3J. Error bars indicate S.D. (n = 3, ** p < 0.01 by Student’s two-tailed unpaired t test).

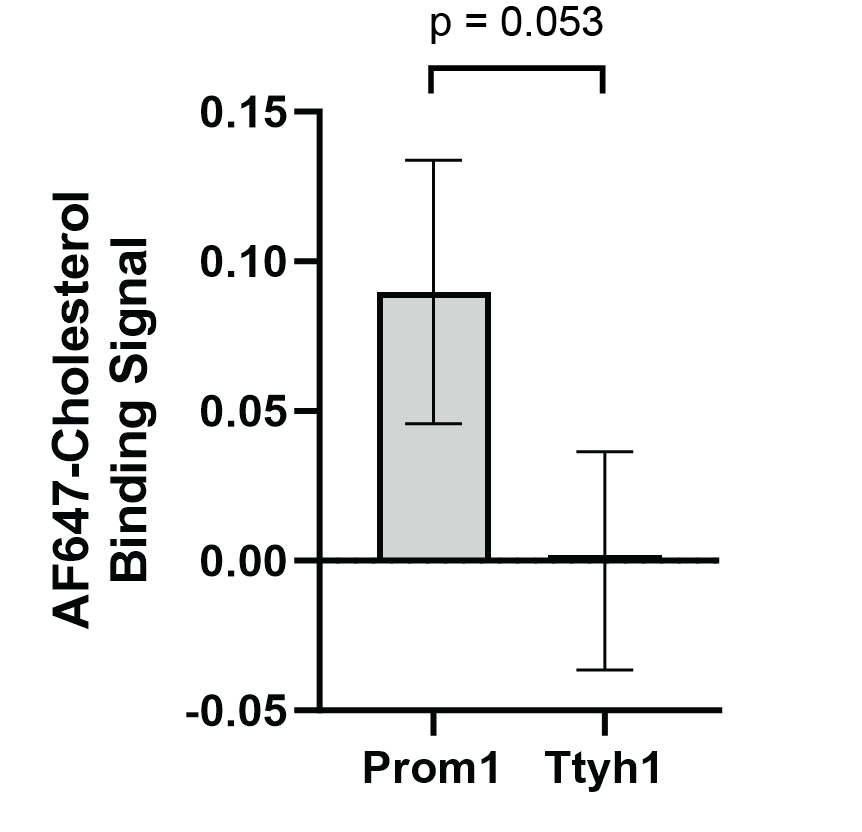

**Figure S7.** Cholesterol co-immunopurification (Chol-IP) measurement of AlexaFluor647-cholesterol binding in Prom1-Strep or Ttyh1-Strep EVs. Error bars indicate S.D. (n = 3, p = 0.053 by Student’s two-tailed unpaired t test).

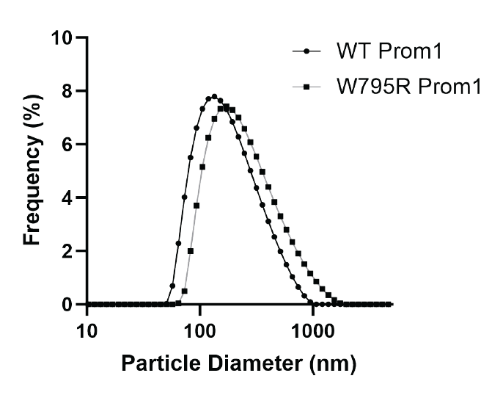

**Figure S8.** Dynamic light scattering EV diameter measurements of purified WT or W795R Prom1-Strep EVs.

| **Prom1 Mutant** | **Specific Mutations** |
| --- | --- |
| Wild-type (WT) | - - |
| CRAC-1 | V460A, Y463L |
| CRAC-2 | Y819L |
| CRAC-3 | L125A, F130L |
| CRAC-4 | L806A, F811L |
| CARC-1 | F158L, L162A, L163A |
| CARC-2 | F804L, L806A, L809A |
| F62L | F62L |
| F505L | F505L |
| F794L | F794L |
| W795R | W795R |
| F796L | F796L |

**Table ST1.** Prom1 mutants used in this study. All mutants are adapted from isoform S1 (NCBI accession NP_001139319.1).

**Supporting Information References**

1.	Clamp, M., Cuff, J., Searle, S.M., and Barton, G.J. (2004). The Jalview Java alignment editor. Bioinformatics *20*, 426–427. 10.1093/bioinformatics/btg430.

2.	Jumper, J., Evans, R., Pritzel, A., Green, T., Figurnov, M., Ronneberger, O., Tunyasuvunakool, K., Bates, R., Žídek, A., Potapenko, A., et al. (2021). Highly accurate protein structure prediction with AlphaFold. Nature *596*, 583–589. 10.1038/s41586-021-03819-2.

3.	Anisimova, M., and Gascuel, O. (2006). Approximate Likelihood-Ratio Test for Branches: A Fast, Accurate, and Powerful Alternative. Syst. Biol. *55*, 539–552. 10.1080/10635150600755453.

4.	Vaughan, T.G. (2017). IcyTree: rapid browser-based visualization for phylogenetic trees and networks. Bioinformatics *33*, 2392–2394. 10.1093/bioinformatics/btx155.
